# Deficient cell-cell cohesion is linked with lobular localization in simplified models of lobular carcinoma *in situ* (LCIS)

**DOI:** 10.1101/2024.12.12.628158

**Authors:** Matthias Christgen, Rodrigo A. Caetano, Michael Eisenburger, Arne Traulsen, Philipp M. Altrock

**Author notes:** MC and PMA contributed equally. Correspondence to: Prof. Dr. med. Matthias Christgen, PhD Institute of Pathology, Hannover Medical School Carl-Neuberg-Str. 1, 30625 Hannover, Germany Philipp M. Altrock, PHD Department of Theoretical Biology, Max Planck Institute for Evolutionary Biology August-Thienemann-Str. 2, 24306 Plön, Germany.

## Abstract

Lobular carcinoma *in situ* (LCIS) is a precursor of invasive lobular carcinoma of the breast. LCIS cells lack cell-cell cohesion due to the loss of E-cadherin. LCIS cells grow in mammary lobules rather than in ducts. The etiology of this pattern, especially its dependence on cellular cohesion, is incompletely understood. We simulated passive intra-glandular scattering of carcinoma *in situ* (CIS) cells in an ultra-simplified hollow mold tissue replica (HMTR) and a discrete-time mathematical model featuring particles of variable sizes representing single cells (LCIS-like particles) or groups of cohesive carcinoma cells (DCIS-like particles). The HMTR features structures reminiscent of a mammary duct with associated lobules. The discrete mathematical model characterizes spatial redistribution over time and includes transition probabilities between ductal or lobular localizations. Redistribution of particles converged toward an equilibrium depending on particle size. Strikingly, equilibrium proportions depended on particle properties, which we also confirm in a continuous-time mathematical model that considers controlling lobular properties such as crowding. Particles of increasing size, representing CIS cells with proficient cohesion, showed increasingly higher equilibrium ductal proportions. Our investigations represent two conceptual abstractions implying a link between loss of cell-cell cohesion and lobular localization of LCIS, which provide a much-needed logical foundation for studying the connections between collective cell behavior and cancer development in breast tissues. In light of the findings from our simplified modeling approach, we discuss multiple avenues for near-future research that can address and evaluate the redistribution hypothesis mathematically and empirically.

## 1. Introduction

Lobular carcinoma *in situ* (LCIS) is a non-obligate precursor of invasive lobular carcinoma (ILC) of the breast (1–3). Loss of E-cadherin is primarily due to mutational inactivation of the *CDH1*/E-cadherin gene (4,5). In histologic sections, LCIS cells resemble “marbles in a bag” (6). Contrary to ductal carcinoma *in situ* (DCIS), LCIS cells grow in the acini of mammary lobules rather than in mammary ducts and form multifocal skip lesions (2,7–10). The etiology of this distribution pattern is incompletely understood.

According to a popular concept developed by Foote and Stewart in 1941, DCIS and LCIS arise from the mammary epithelium of different anatomic compartments, namely ducts and lobules (11). This was widely accepted as an explanation for the differences between DCIS and LCIS. Multifocality of LCIS was viewed as independent tumor initiation in multiple lobules (11). Histological studies by Wellings *et al.* in 1975 indicated that DCIS and LCIS both arise in terminal ducto-lobular units (TDLU, i.e., lobules and associated terminal ducts) (12). On the one hand, Wellings *et al.* provided histologic evidence against different anatomic origins of DCIS and LCIS. On the other hand, this study could not explain the preferential growth of LCIS in lobules or its multifocality. Molecular analyses identified E-cadherin loss and mutation of *CDH1* in ILC and LCIS as important events (4,5). Sakr *et al.* and others employed mutational profiling to address clonal relatedness in multifocal LCIS (13,14). Strikingly, multifocal LCIS lesions often harbor a single, unique somatic *CDH1* mutation (13,14). This demonstrates a shared genetic ancestry in multifocal LCIS and argues against the concept of independent tumor initiation in multiple lobules, as envisioned earlier (13,14).

An alternative concept for the spatial development of LCIS involves the passive intra-glandular scattering of LCIS cells through mammary ducts (13,14), which may be termed the “redistribution concept.” Tumor initiation may occur anywhere in the epithelium. At the initiation site, LCIS cells shed off from the transformed epithelium. Subsequently, LCIS cells scatter through mammary ducts and colonize secondary intra-glandular sites. This theoretical concept would harmonize the histologic feature of multifocality (skip lesions) with the empirical finding of shared *CDH1* mutations in multifocal LCIS lesions (13,14).

A shortcoming of this theoretical redistribution concept has been that it does not explain the preferential growth of LCIS cells in lobules instead of ducts. One possible explanation is that loss of cell-cell cohesion provides LCIS cells with a high capability for lobular colonization due to a higher probability of entering a lobule and getting trapped. Compared to DCIS cells, which form cohesive aggregates, single non-cohesive LCIS cells may have a greater probability of entering the narrow acini of mammary lobules. Once arrived at this micro-environmental niche, mammary lobules may trap scattered LCIS cells in their complex structure of branching acini (15). Subsequently, LCIS cells may replace normal luminal epithelial cells of affected acini.

Studying the spatial development of LCIS is hampered by the lack of longitudinal *in vivo* models that allow the monitoring of dynamic cell distribution patterns on a microscopic (cellular) or mesoscopic (small cell population) level (16,17). Further, anatomical differences exist between the human breast and the mammary gland of common laboratory animals, such as mice. The human breast develops permanent lobules at puberty, which enlarge during pregnancy but do not undergo complete involution until the senium (18,19). The mouse mammary gland develops lobules during pregnancy, which later regresses almost completely, complicating the modeling of LCIS in mice (20,21).

Mathematical models of biological systems and diseases, such as cancer, can integrate mechanistic assumptions with prior knowledge and new data. Like experimental model systems, mathematical models of biological systems span multiple scales. Models of cell populations are of particular significance for the dynamics of complex tissue phenomena, e.g., anti-cancer treatment, cancer evolution, and the biophysical processes of tissue confinement and cellular movement (22–26). These theoretical models can quantify empirically tested or proposed mechanisms. They allow us to abstract biological phenomena, explore the influence of mechanistic or technical assumptions on a system’s kinetics or dynamics, infer model parameters, and enable validation or testing of models in light of available experimental or clinical data (27,28).

In this study, we model passive intra-glandular scattering of carcinoma *in situ* (CIS) cells. We used crowds of tiny steel beads moving in an ultra-simplified hollow mold tissue replica (HMTR) featuring one mammary duct connected to several mammary lobules and mathematical descriptions of this dynamical system. We used a discrete-time mathematical model to precisely match the HTMR dynamics and a more general continuous-time (ODE) model to investigate the theoretical aspects of the lobular redistribution concept under the influence of crowding as a possible result of particle size dependence. Our findings suggest a potential mechanistic link between the loss of cell-cell cohesion and the predominantly lobular localization of LCIS.

## 2. Materials and Methods

### 2.1 Design of the hollow mold tissue replica (HMTR)

Based on morphometric measurements in normal human mammary gland tissue (**Figure 1** and **Supplemental Data Table 1**), we devised an ultra-simplified, two-dimensional hollow mold tissue replica (HMTR) at a scale of 80:1 (**Figure 2A**). The HMTR featured one curled duct (6.0 mm width) and six lobules (40.0 mm size). Lobules were designed as small ductal ramifications with three dichotomous branching generations and eight acini (approximately 2.0 to 2.5 mm width) each (**Figure 2B**). This aligns with the microscopic anatomy, but original lobules are more complex (up to twelve dichotomous branching generations) (15). All anatomical structures (duct, ducto-lobular connection segments, lobules, acini) were milled in an ordinary plywood board (280.0 x 140.0 x 4.0 mm) using a Dremel 4000 precision rotary tool (Dremel, Breda, The Netherlands) equipped with a 2.0 mm high-speed cutter (Dremel). The resulting HM plate was sealed with acrylic varnish and glaze (AquaClou, Hornbach, Bornheim, Germany). The HM plate was sandwiched between a transparent cover plate (280.0 x 110.0 x 2.0 mm, polysterol, Palram, Schönebeck, Germany) and a solid floor plate (280.0 x 140.0 x 4.0 mm, plywood) using machine screws (M3 x 16.0 mm) and butterfly nuts (M3). A grid (10.0 x 10.0 mm) was printed on the cover plate to improve display (**Figure 2C**). For wobble shaking, the HMTR was attached to a rotary table (280.0 x 180.0 x 6.0 mm, plywood) and pegged on a rotary axle (220.0 x 30.0 mm, beech wood). A plywood casing consisting of two lateral plates (300.0 x 300.0 x 6.0 mm, plywood) and four horizontal corner pillars (188.0 x 35.0 x 35.0 mm, spruce wood) served as an axle bearing. The plywood casing also featured a handle for the rotary axis and striker pins to adjust the wobble-shaking angle (**Figure 2A**). A detailed constructional drawing is provided in the supplement (**Supplemental Data Figure 1**).

**Figure 1.**
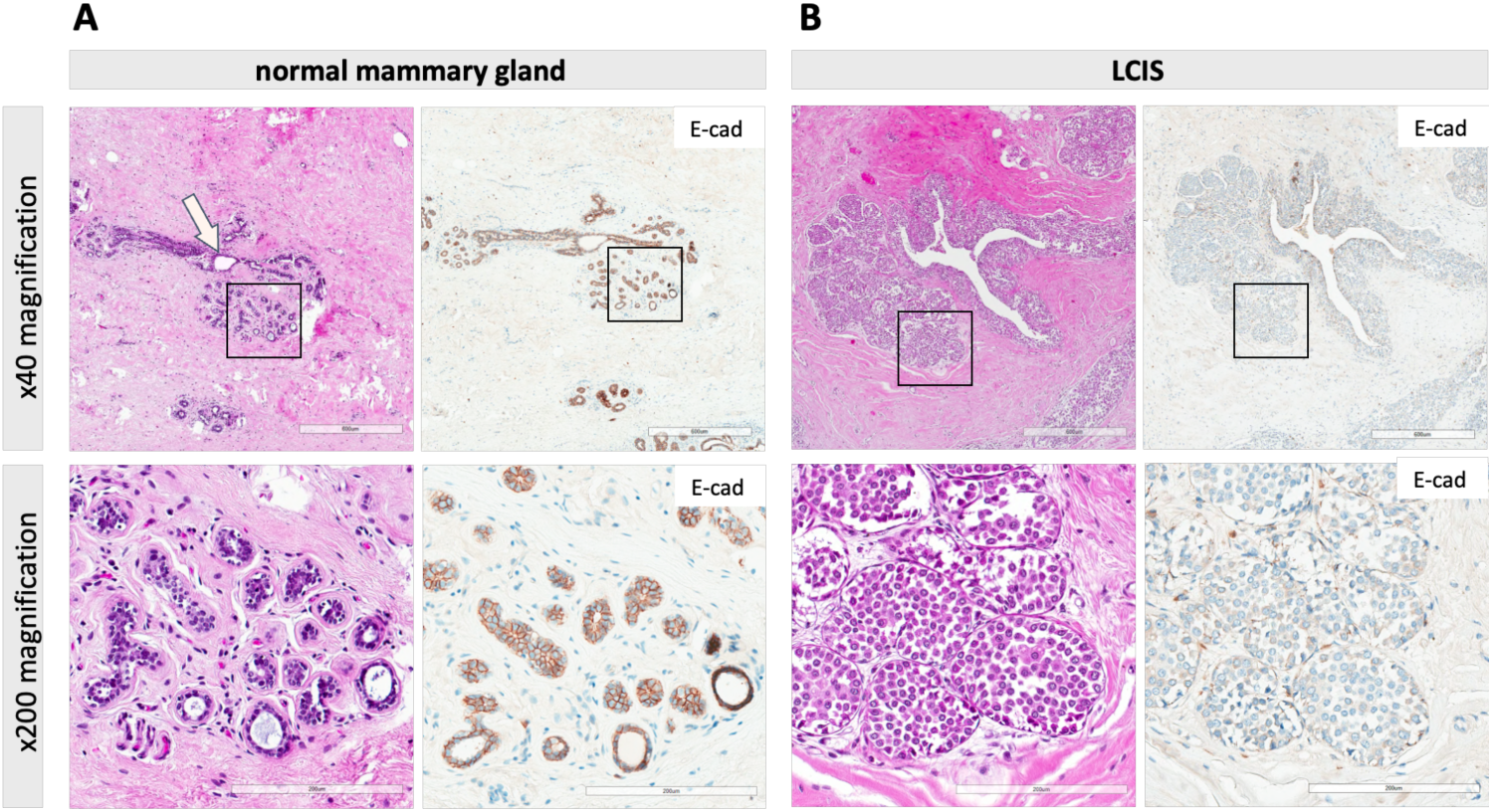
Histology of the normal mammary gland and LCIS. **(A)** Representative photomicrographs of HE-stained sections of normal mammary gland tissue at x40 magnification (upper panel) and at x200 magnification (lower panel). Scale bars correspond to 600 µm (upper panel) or 200 µm (lower panel). Photomicrographs of IHC stainings for E-cadherin (antibody clone ECH-6) are also provided (right panels). Please note that the depicted lobule has a total size of approximately 750 µm, the duct (white arrow) has a maximum width of approximately 150 µm, and the acini have a width of up to 35 µm. These sizes are consistent with systematic morphometric measurements reported in Supplemental Data Table 1. **(B)** Representative photomicrographs of LCIS at x40 magnification (upper panel) and at x200 magnification (lower panel). Please note the loss of E-cadherin expression (right panel) and the lack of cell-cell cohesion, which results in the characteristic aspect known as “marbles in a bag.”

**Figure 2.**
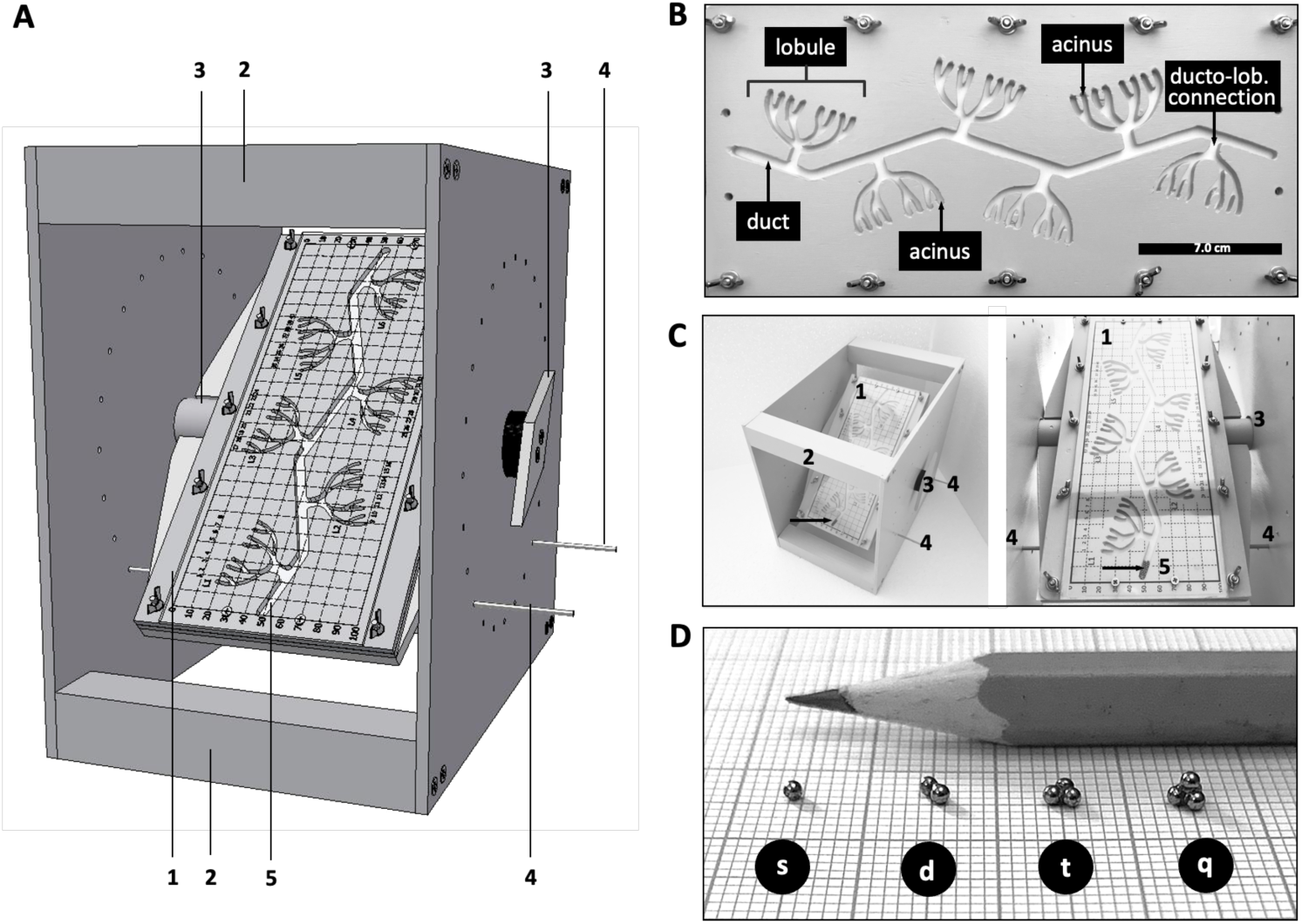
The HMTR of the mammary gland. **(A)** Schematic overview. Parts are as follows: 1, hollow mold (HM) plate with a floor plate and transparent cover plate attached to the rotary table; 2, corner pillar of the casing; 3, handle of the rotary axle; 4, striker pin inserted into the lateral plates of the casing (striker pins adjust the maximum shaking angle of the rotary table to 45 degrees); 5, duct milled into the HM plate. **(B)** Detailed view of the HM tissue replica (HMTR) showing one curled duct (6.0 mm width), six ducto-lobular connections, six lobules, and a total of forty-eight lobular acini (2.0 mm width). **(C)** Photograph of the HMTR fitted in the casing. Please note the charge in the duct (the arrow points at n=24 single beads in the duct). **(D)** Photograph of the different particle types (s, single beads; d, doublet; t, triplet; q, quartet) shown on a 1.0 x 1.0 mm grid. The pencil in the background illustrates the tiny size of the steel beads.

### 2.2 Steel beads used as dummy carcinoma *in situ* (CIS) cells

The HMTR was charged with crowds of steel beads representing motile CIS cells (1.5 mm diameter, GAP ball bearings, Enger, Germany). Bead size (1.5 mm) and duct width (6.0 mm) provided a true-to-scale size ratio (corresponding to CIS cells of approximately 19 µm diameter in a duct of approximately 75 µm width). Different particle types included: i) single beads (to model deficient cell-cell cohesion) and groups of beads glued together as composite beads, namely: ii) doublets, iii) triplets and iv) quartets (to model DCIS-type of CIS cells with gradually more proficient cell-cell cohesion, as reflected by increasing bonds per bead) (**Figure 2D**). Different HMTR charges included: i) N=24 single beads, ii) N=12 doublets, iii) N=8 triplets, or iv) N=6 quartets (**Table 1**). For composite beads (doublets, triplets, and quartets), we used single beads glued with a special adhesive (Prime&Bond active^®^, Densply Sirona, Konstanz, Germany).

**Table 1.** Particle types and charges.

| particle type | bonds per bead | particle size | charge |
| --- | --- | --- | --- |
| single | 0 | 1.5 x 1.5 x 1.5 mm | 24 |
| doublet | 1 | 3.0 x 1.5 x 1.5 mm | 12 |
| triplet | 2 | 3.0 x 2.8 x 1.5 mm | 8 |
| quartet | 3 | 3.0 x 2.8 x 2.8 mm | 6 |

Overview of particle types. Charge refers to the number of particles that were initially placed in the duct; the number of beads was conserved, but as a consequence, the number of particles decreased with increasing bonds per bead.

### 2.3 Monitoring of redistribution dynamics

Bead or particle scattering was achieved by gentle, manual wobble shaking, as illustrated in **Supplemental Data Figure 2**. The rotary table was rested on the left striker pin (45-degree inclination to the left side). Next, beads were placed in the left end of the duct, and the HMTR was covered with a transparent cover plate to prevent the loss of particles. Next, the rotary axle was gently turned clockwise until the rotary table touched the right striker pin (45-degree inclination to the right side; redistribution cycle 1). Due to this motion, particles moved down the duct from left to right, and some particles entered any of the six lobules (**Supplemental Data Figure 2**, red arrows). The number of particles located in the duct, in each ducto-lobular connection segment, and in every lobule was recorded after each axle rotation (after each redistribution cycle). Each particle type (single beads, doublets, triplets, and quartets) was monitored for n=20 redistribution cycles (reflecting elapsed time). Single beads were scored as single events, while doublets, triplets, and quartets were scored as two, three, and four events, respectively, to preserve the total number of beads. All experiments were repeated four times (2x start position on the left, 2x start position on the right side of the HMTR). Accordingly, bead/particle distribution patterns were recorded over 336 time points (4 bead types x 20 cycles x 4 experimental replicates = 320; + 16 time points for each initial condition = 336).

### 2.4 Discrete mathematical model of bead redistribution dynamics

To approximate the dynamics of the HTMR experiment in a more abstract, potentially generalizable way, we devised a mathematical model of the bead or particle redistribution dynamics. We represent the system by two states: a duct *D,* connected to *M* lobule-like regions that can trap the particles, *L_i_*, whereby the integer *i* ranges from 1 to *M.* Note that *M*=6 in the abovementioned HTMR experiments. Every particle can thus be in state *D* or state *L_i_*. Particles can move between *D* and the *L_i_*. To define the state *L,* we took the sum over all *L_i_*. Time is discrete, mimicking the experimental shaking dynamics in discrete cycles. Within one time step, particles in the duct (in state *D*) can stay in *D* or enter one of the lobules *L_i_*. We assume that for each particle, upon passing the opening to a lobule, the probability of getting into a lobule is *p*_in_. Similarly, beads in state *L* (i.e., in any of the *L_i_*) can either move to state *D* with probability *p*_in_or stay in *L* with probability 1 – *p*_in_ (illustrated in **Figure 3A)**.

**Figure 3.**
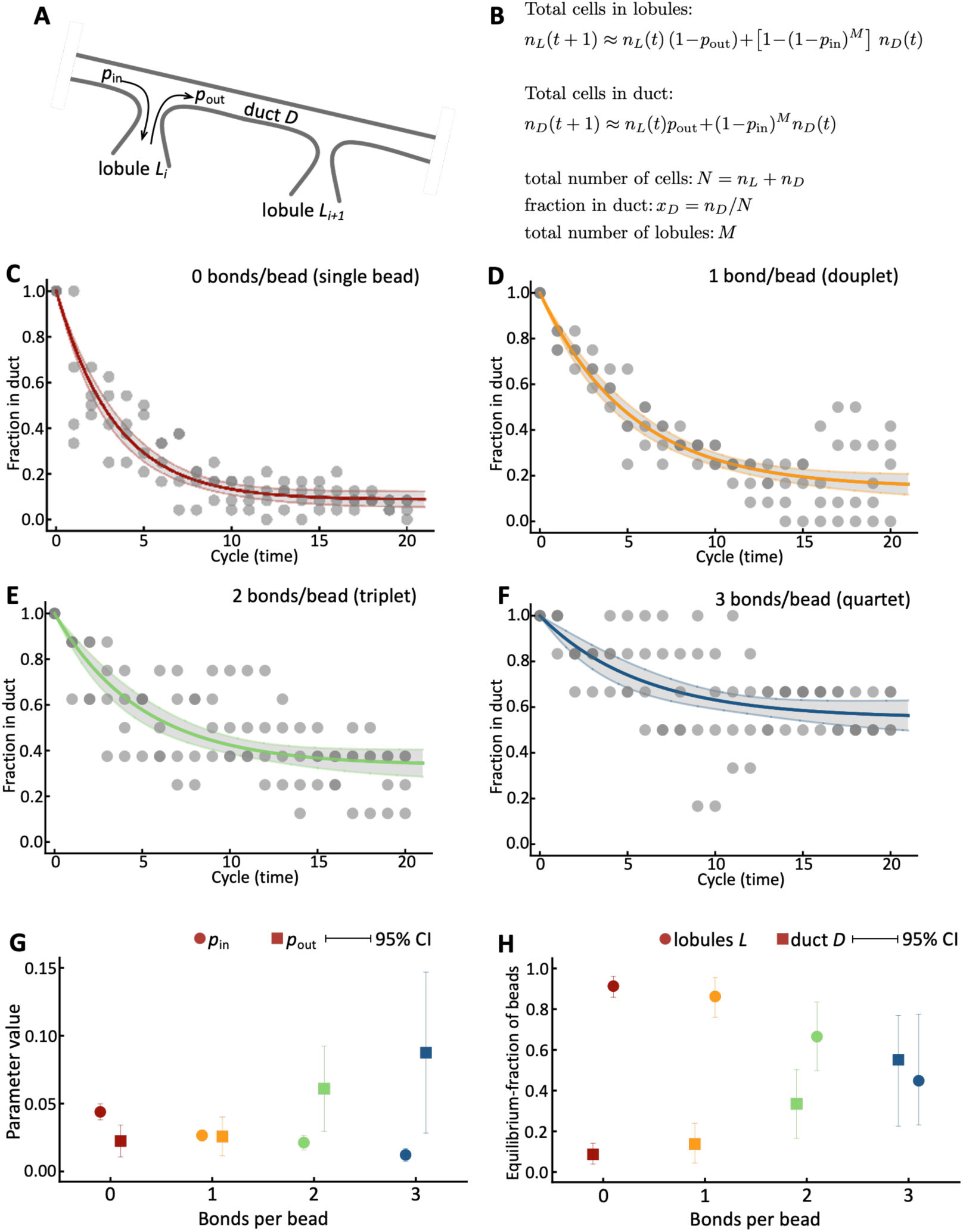
Discrete mathematical model and data fitting. **(A)** Schematic of the kinetic mathematical model. Beads or particles can move from duct, state *D*, into a lobule, states *L_i_* (there are *i*=1,2,…,*M* lobules) with probability *p*_in_. Beads can move from a lobule to the duct with a probability *p*_out_. *M*=6 in all following experiments. **(B)** We derived a mathematical model for the dynamics of particle redistribution (Methods). **(C)** The model fits experimental data using least squares minimization. Shown are data from four experimental replicates (dots), the best fit, and the 95% confidence interval of the model (see methods section) for single beads (zero bonds per bead) all initiated in the duct. **(D)** Model fit to data obtained with doublets (particles of two beads, one bond per bead). **(E)** Model fit to data obtained with triplets (particles of three beads, two bonds per bead). **(F)** Model fit to data obtained with quartets (particles of four beads, three bonds per bead). **(G)** The inferred parameters *p*_in_ and *p*_out_ for the four particle types (symbols: best-fit values; whiskers: 95% CI). **(H)** Best-fit equilibrium particle distributions as a function of bonds per bead.

We are interested in the dynamics of particles, which can be single beads, doublets, triplets, or quartets. Let us label the particles in state *D* as *n_D_*(t), and those in the states *L_i_* as *n_L_*_,*i*_(t). The total number of particles, *N,* is conserved: *n_D_*(t)+*n_L_*_,*1*_(t)+*n_L_*_,*2*_(t)+…+*n_L_*_,*M*_(t)=*N*. Under the assumption of a sequential arrangement of lobules, we can write a discrete-time master equation for the number of particles the *L_i_*:

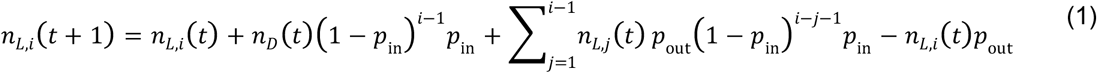

This equation states that the number of particles in *L_i_* after a single time step (on the left-hand side) is equal to the particles that were already there (first term on the right-hand side), plus the particles that are gained from *D* (second term), plus the ones that are gained from other lobules (the sum, third term), minus the particles that leave (fourth term). To end up in lobule *i*, a particle in *D* must pass all previous lobules, which occurs with probability (1 – *p*_in_)*^i^*^-1^. In the experimental setup, only a subset of lobules can contribute to gain in lobule *i*, which occurs with probability *p*_out_(1 – *p*_in_)^i-j-1^—to have a chance to enter lobule *i*, a particle must first leave lobule *j* and then not enter any lobules in between.

Similarly, we can formulate a master equation for the number of beads in the duct:

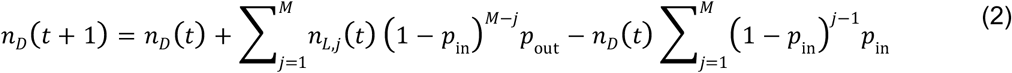

Here, the gain from a particular lobule (second term on the right-hand side) and the loss term (third term) must account for the condition that the particles are not lost to other lobules. The last term in Eq. (2) can be simplified

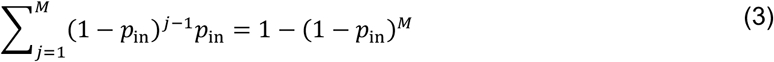

This system of *M*+1 equations given by formulas (1) and (2) describes the dynamics of particles in the HTMR setting. We can assume that *p*_in_ and *p*_out_ are small to simplify the system further, i.e., we can neglect terms with products such as *p*_in*_*p*_out_, etc. This allows us to sum over all lobule compartments 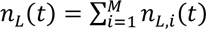, and we can approximately describe the dynamics by two equations:

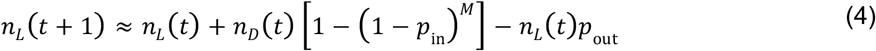

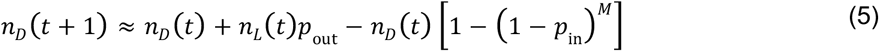

This system can be solved recursively. Noting that *n_D_*(*t*) + *n_L_*(*t*) = *N* is constant, and assuming that all particles are in the duct initially, *n_D_*(0) = *N*, we obtain a solution for the number of particles in *D* over time

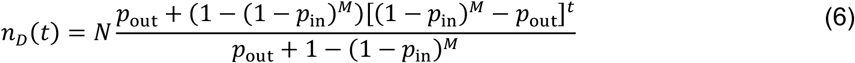

and the number of particles in the lobules follows as *n_D_*(*t*) = *N* – *n_L_*(*t*). This solution only depends on the probabilities of moving in and out of a lobule, the number of lobules, and the number of particles. For a large enough number of lobules, *M*, [(1 – *p*_in_)^8^ – *p*_out_]*^t^* is positive. This factor will converge to 0 as time becomes very large, *t* → ∞, and the system equilibrates. The equilibrium number of particles in the duct is thus given by the time-independent formula

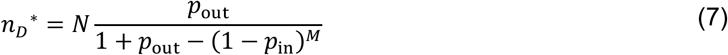

Equation (6) can be used to infer the probabilities *p*_in_ and *p*_out_for a given particle size (1 bead, 2 beads, etc.) based on longitudinal observations. Equation (7) can be used to calculate the long-term proportions of particles in the duct *n_D_*^∗^⁄*N*, and in the lobules 1 – *n_D_*^∗^⁄*N*.

### 2.5 Parameter estimation

The bead or particle redistribution dynamics were fitted with least-squares minimization (*NonlinearModelFit*, Wolfram Mathematica, version 13.3), using Eq. (6), and the longitudinal experimental data. The proportion of particles in the ductal or the lobular compartments was used as input data, and Eq. (6) was evaluated at times *t*=0,1,2,… using *n_D_*(*t*)/*N*, minimizing the joint sum of squares to four replicate experimental time series. The total number of particles *N* was fixed (*N*=24 for single beads, *N*=12 for doublets with one bond per bead, *N*=8 for triplets with two bonds per bead, and *N*=6 for quartets with three bonds per bead). Each experiment was conducted with *M*=6 identical lobules. From the fitting procedure, best-fit values for *p*_in_ and *p*_out_ were obtained, with the 95% confidence intervals calculated using the standard estimate of the error, shown in **Table 2**.

**Table 2.** Mathematical model fit results.

| parameter* | particle type | best fit value<sup>§</sup> | 95CI |  |
| --- | --- | --- | --- | --- |
|  |  |  | lower | upper |
| $p_{in}$ | singles | 0.0439 | 0.0379 | 0.0499 |
|  | doublets | 0.0256 | 0.0226 | 0.0306 |
|  | triplets | 0.0213 | 0.0159 | 0.0267 |
|  | quartet | 0.0122 | 0.0075 | 0.0169 |
| $p_{out}$ | singles | 0.0225 | 0.0108 | 0.0341 |
|  | doublets | 0.0230 | 0.0115 | 0.0401 |
|  | triplets | 0.0610 | 0.0297 | 0.0924 |
|  | quartet | 0.0875 | 0.0282 | 0.1468 |

\* $p_{in}$ : probability of getting from the duct ( $D$ ) into any lobule ( $L$ ); $p_{out}$ : probability to move out of a lobule ( $L$ ) into the duct ( $D$ ).
<sup>§</sup>Best fit parameters refer of the mathematical model (Eq. (6)) for the four different particle types. Fits were performed using the fraction of particles in the duct, with $M=6$ . 95CI: 95% confidence interval.

## Results

### 3.1 A hollow mold tissue replica (HMTR) to model CIS cell scattering

In histological sections, LCIS cells display a characteristic aspect known as “marbles in a bag” (**Figure 1**) (6). This aspect is due to the loss of cell-cell cohesion (6,9). The metaphor refers to the morphologic impression that LCIS cells move around in the lumina of affected mammary gland structures (9). This study proceeded from the theoretical assumption that, to some extent, this passive movement is a real feature of LCIS. Accordingly, we sought to model passive intra-glandular scattering of carcinoma *in situ* (CIS) cells using crowds of tiny steal beads in a hollow mold tissue replica (HMTR) of mammary gland tissue. Based on morphometric measurements (**Supplemental Data Table 1**), we constructed an ultrasimplified, two-dimensional HMTR at a scale of 80:1. The HMTR featured one mammary duct, six lobules, and a total of forty-eight lobular acini (**Figure 2A-C**). Detailed construction drawings are provided in the supplement (**Supplemental Data Figure 1**).

The duct was charged with steel particles to approximate motile CIS cells (**Figure 2D**). Scattering was achieved by gentle wobble shaking at a 45-degree angle. For this purpose, we attached the HMTR to a rotary table fitted in a half-open casing (**Figure 2A**). Different particle types and charges included: i) N=24 single beads (representing CIS cells with deficient cohesion; LCIS-type), or composite-bead particles, namely ii) N=12 doublet beads, or iii) N=8 triplet beads, or iv) N=6 quartet beads, representing groups of CIS cells with gradually more proficient cohesion. This procedure provided increasingly larger particles (**Figure 2D** and **Table 1**). The total number of beads was always constant (total load equivalent to N=24 single beads). Upon wobble shaking, beads scattered in the HMTR and redistribution patterns were monitored for 20 redistribution cycles. The schematics of two representative redistribution cycles are illustrated in **Supplemental Data Figure 2**. A mathematical model was devised to describe the redistribution dynamics toward an equilibrium distribution across the duct and lobules (**Figure 3A, B**) and to estimate the particle size-dependent kinetic parameters of the system.

### 3.2 Single beads were rapidly cleared from the duct

The fraction of beads or particles in the duct decreased with each redistribution cycle, i.e., with elapsed time (**Figure 3C-F**). Single beads (**Figure 3C**) were rapidly cleared from the duct and redistributed to various lobules. For instance, after 5 cycles, approximately 36% of single beads were in the duct, and approximately 64% were in lobules. After 10 cycles, approximately 11% of single beads remained in the duct, and approximately 89% were in lobules (**Figure 3C**).

A mathematical model also described the redistribution dynamics (Methods, **Figure 3B**). The redistribution rate and the approximate equilibrium strongly depended on particle size and thus on particle type. For single beads, a rapid decay toward an equilibrium was observed **(Figure 3C)**. For doublet, triplet, and quartet particles, this equilibration was substantially slower and more widely distributed **(Figure 3D-F)**. Based on our model fits, the probability of a bead moving into a lobule (*p*_in_) decreased with increasing particle size (compare *p*_in_=0.0439 [95CI: 0.0379-0.0499] for single beads to *p*_in_=0.0122 [95CI: 0.0075-0.0169] for quartets) **(Table 2)**. Similarly, the probability of particles moving from a lobule to the duct increased (compare *p*_out_ =0.0225 [95CI: 0.0108-0.0341] for single beads to *p*_out_=0.0875 [95CI: 0.0282-0.1468] for quartets) **(Table 2)**. The change in inferred redistribution probabilities (*p*_in_ and *p*_out_) is shown in **Figure 3G**. Interestingly, for doublet particles, we estimated that the probabilities were roughly equal, *p*_in_=*p*_out_, whereas, for quartet beads, the probability of moving from a lobule to the duct was approximately 7-fold higher than the probability of moving from the duct into a lobule (*p*_in_<*p*_out_) **(Figure 3G)**.

### 3.3 Composite bead particles were redistributed slowly and more likely remained in the duct

Composite bead particles (doublet, triplet, and quartet beads), representing groups of CIS cells with gradually more proficient cell-cell cohesion, were cleared from the duct much slower than single beads (**Figure 3C-F**). These particles were also characterized by drastic changes in the kinetic parameters *p*_in_ and *p*_out_ (**Figure 3G**). Composite bead particles reached equilibrium distributions at increasingly higher fractions in the duct (**Figure 3H**). Single beads, doublets, triplets, and quartets plateaued at 8.7% [95CI: 3.9-14.2%], 13.8% [95CI: 4.4-23.9%], 33.5% [95CI: 16.5-50.2%], and 55.1% [95CI: 22.5-76.9%]) intraductal fractions, respectively (**Table 3**). Smaller fractions were eventually located in the lobular compartment (**Figure 3H**). The mathematical model could explain these results based on the kinetic parameters *p*_in_ and *p*_out_. While *p*_in_ decreased and *p*_out_ increased with increasing particle size, they did so at different rates. The ratio *p*_out_/*p*_in_ was approximately 0.5 for single beads, approximately 1.0 for doublets, 2.9 for triplets, and >7.0 for quartets. Single beads were 2-fold more likely to move into a lobule than out; doublets moved in and out with equal likelihoods. The largest particles, namely quartet beads, were 7-fold more likely to move out of lobules.

**Table 3.** Equilibrium particle distributions based on mathematical model fits.

| bead type | equilibrium | equilibrium | 95CI for $x_D$ | |
| --- | --- | --- | --- | --- |
| | fraction in $L$ | fraction in $D$, $x_D$ | lower | upper |
| singles | 0.913 | 0.087 | 0.0392 | 0.1415 |
| doublets | 0.862 | 0.138 | 0.0444 | 0.2392 |
| triplets | 0.665 | 0.335 | 0.1654 | 0.5019 |
| quartets | 0.449 | 0.551 | 0.2249 | 0.7687 |

### 3.4 Many-lobule systems dynamics and lobular crowding

We can devise a slightly more general dynamical system of a duct and multiple lobules to explore possible mechanisms of particle interactions in lobules. Let us now assume that during a short interval *dt*, a cell or group of cells (“particle”) in the duct can move into a given lobule at a rate that depends on the state of the lobule *q_i_* and out of the lobule at rate *p_i_*. Let *x_d_* be the number of particles in the duct, and *x_i_* the number of particles in lobule *i*. Then, we can describe the dynamics between these compartments with the following ordinary differential equations that keep the initial density of particles constant (*dx_d_*⁄*dt* + ∑_*i*_ *dx_i_*⁄*dt* = 0)

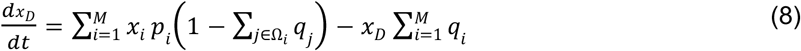

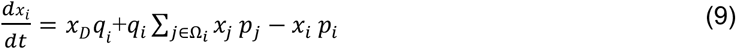

Here, Ω_3_ is the set of lobules that can scatter into lobule *i* in the time interval *dt*. For example, nearest-neighbor interactions lead to Ω_3_ = {*i* − 1, *i* + 2} with the special cases Ω_5_ = {2} and Ω_*M*_ = {*M* − 1}. We can model the filling of lobules, or crowding, by imposing a constant *c* and the sensitivity parameter β:

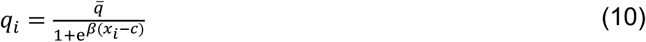

This function reduces the rate of particles moving into a lobule, down from the baseline value 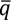. This model is a phenomenological approach to describe how lobular crowding might affect the distribution of cells or cell clusters across duct and lobules. We explore the continuous model, Eqs. (8) and (9), in **Figure 4**. Analogously to the discrete model, the fraction of particles in the duct assumes an equilibrium value (**Figure 4A**), and the equilibrium fraction in the duct decreases with the number of lobular compartments, *M* (**Figure 4B**). We considered a monotonously decreasing rate of moving from duct into a lobule (Eq. (10)), examples of which are shown in (**Figures 4C,D,E**). This rate function is by where it assumes its half-maximal value, *c*, and by its maximal slope i.e., the intensity parameter β (**Figure 4E**). With increasing values of c, the equilibrium fraction in the duct decreases (**Figure 4F**). The sensitivity β can have opposing effects, depending on the value of *c* (**Figure 4G**). If *c* is sufficiently large, counterintuitively, more particles are expected to get trapped in lobules because the detrimental impact of crowding only takes effect after sufficiently many particles have been trapped. Thus, depending on the lobular potential to trap particles, fewer (low *c*) or more (high *c*) particles can get trapped due to crowding. High threshold values of the rate to enter lobules, possibly due to crowding, decrease the duct fraction nonlinearly. This effect depends on the number of particles available to overcrowd lobules. For sufficiently low thresholds (values of *c*), the fraction in the duct could be higher than in the absence of crowding.

**Figure 4.**
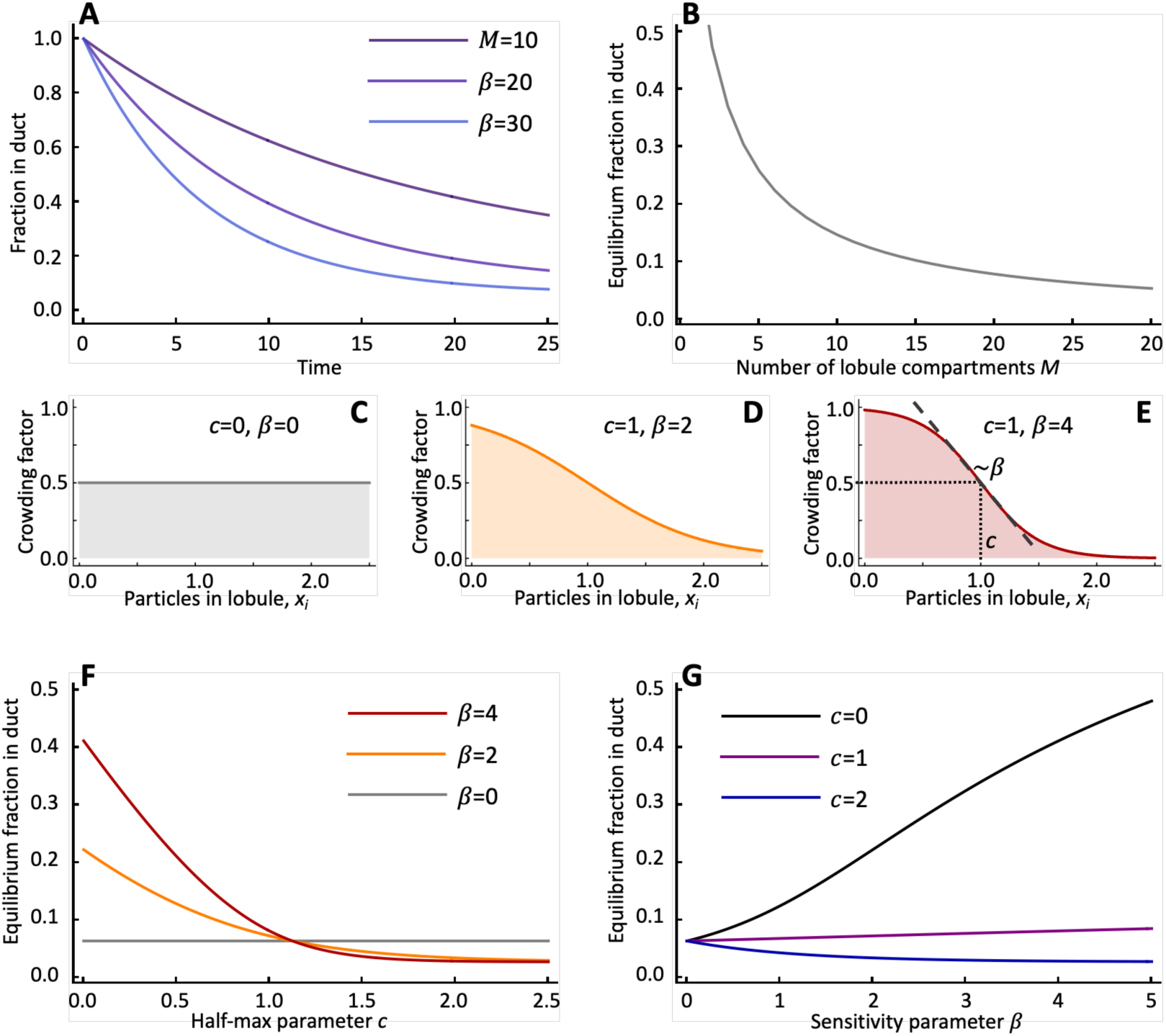
A continuous mathematical model to investigate lobular crowding, Eqs. (8) and (9). **(A)** Temporal evolution of the fraction in the duct, *x_D_*(*t*)⁄*x_D_*(0), for *p_i_* = *q_i_* = 0.01. **(B)** Decline of the fraction in the duct, *p_i_* = *q_i_* = 0.1. **(C)**, **(D)**, **(E)** The behavior of *q_i_*, Eq. (10), for various parameter values, 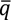. **(F)** Change of ductal fraction with increasing capacity of each lobule. **(G)** Change of ductal fraction with increasing sensitivity to crowding. In all panels, the initial conditions were 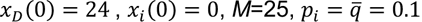, unless specified otherwise.

## 4. Discussion

LCIS cells typically lack cell-cell cohesion due to E-cadherin loss and grow in mammary lobules rather than in mammary ducts (8–10). In histological sections, LCIS cells display a characteristic aspect known as “marbles in a bag” (6,9). This metaphor refers to the morphological impression that LCIS cells move around in the lumina of mammary gland structures. The development of LCIS is incompletely understood, especially concerning distribution patterns, multifocality, and preferential lobular localization. Multifocality has traditionally been explained by synchronous, independent tumor initiation in multiple lobules (11). However, recent genetic evidence indicates that multifocal LCIS lesions share the same genetic ancestry, which argues against independent tumor initiation at multiple sites (14,29).

The redistribution concept is one of several alternative theoretical concepts of LCIS development that could harmonize multifocality and clonal relatedness. According to the redistribution concept, LCIS development involves the passive scattering of clonal CIS cells through mammary ducts and subsequent colonization of various mammary lobules. Compared with cohesive aggregates of DCIS cells, single non-cohesive LCIS cells may have a greater probability of entering the narrow acini of mammary lobules. To illustrate the redistribution concept, we modeled passive intra-glandular scattering of carcinoma *in situ* (CIS) cells with crowds of tiny steal beads in an HMTR. The HMTR was devised explicitly for this experiment and featured one mammary duct and six lobules. Size dimensions were based on histologic measurements in actual human breast tissue, up-scaled in a ratio of 80:1. Passive bead scattering was achieved by wobble shaking, and bead redistribution from the ductal to the lobular compartment was also described in a kinetic mathematical model. This mathematical model is semi-mechanistic as it does not specify how the probabilities of moving in and out of a lobule are a function of size or shape, but provides a valuable foundation for more sophisticated computational modeling and stimulates further research.

Our results show that single beads (representing CIS cells with deficient cohesion; LCIS-type) were rapidly cleared from the duct and redistributed to various lobules. Under otherwise constant experimental conditions, larger composite bead particles (representing CIS cells with proficient cohesion; DCIS-type) showed a different behavior: larger particles tended to remain in the duct. The number of bonds per bead (representing the degree of CIS cell cohesiveness) and the equilibrium proportion of intra-ductal remainders were linked, as evidenced by a nearly linear relation (**Figure 3H**). Hence, our experimental HMTR model and the mathematical modeling suggest a potential mechanistic link between the loss of cell-cell cohesion and the predominantly lobular localization of LCIS. In agreement with experimental findings, the discrete mathematical model (formulated to match the experimental setting) found that the kinetics of moving from lobules into the duct increased with the number of bonds. A more general continuous-time mathematical model confirmed this finding and could be used to describe possible lobular crowding effects that would alter the expected fraction of particles in the duct, possibly in a nonlinear fashion.

Our study has numerous limitations. The HMTR is a non-physiological, mechanical model. Neoplastic cells are not steel beads. Real mammary lobules are highly complex, three-dimensional (30,31), and flexible structures (15) that may alter the microscopic movement of cells. Often, in developmental and cancer biology, such movement-induced changes in the density of cells are captured in reaction-diffusion-advection systems (32) that, in principle, allow numerical or even approximate analytical solutions. Agent-based models present a more computationally involved approach in which cells and their surrounding tissue are modeled individually with a set of (complicated) individual rules determining possibly stochastic cellular movements (33,34). These approaches can recapitulate and predict some essential features of growth and invasion, such as self-organization and spatial interactions, including soft-tissue properties (35,36), chemotaxis (37), and the fact that tissues present a low Reynolds number, highly viscous environment that is not considered here. Recent results indicate that extracellular fluid viscosity may enhance cell migration and cancer dissemination (38). Thus, neglecting elevated viscosity in the surrounding medium of cell movement might lead to underestimating the rates of particle movements in and out of lobules and along ducts. Studying the spatial development and redistribution of LCIS is hampered by the lack of longitudinal *in vivo* models that allow the continuous monitoring of dynamic cell distribution patterns because mammary lobules constitute a permanent compartment in the human breast but are a temporary compartment in lab animals (such as mice).

The factors and processes not included in our modeling comprise i) cell migration, diffusion, and chemotaxis, ii) cell death, iii) cell division, iv) tissue flexibility, v) fluctuation of intra-glandular fluids, and vi) cell-cell interactions, such as between CIS cells and the normal luminal epithelium. Most importantly, the singularly directed force of movement leading to redistribution in our experimental approach is likely not the driving force responsible for the passive scattering of CIS cells in actual mammary glands. Fluctuations of intra-glandular fluids due to local tissue pressure changes associated with breathing and walking may be involved in this process.

Another reason for the lack of present-day experimental evidence and the difficulty of designing new longitudinal studies is that regularly formed hollow ductal and lobular gland structures cannot be easily propagated *in vitro*. Few works have described the generation of branching ductal-like structures obtained from 3D cultures of human mammary epithelial cells. These structures show aberrant branching and are not typically hollow (39). Thus, little has been known about many physical micro-environmental conditions in the mammary gland (40–42). Accordingly, how much bead distribution patterns in the HMTR can inform CIS cell distribution in mammary glands is speculative. However, recent work points to essential advances concerning relevant murine-derived organoid models (43). In this context, a challenge will be to couple organoid development with carcinogenesis and to control the site of emergence of abnormally growing cells that move passively or actively.

The HMTR and our mathematical modeling illustrate a theoretical consideration that we summarize under the term “redistribution concept” of LCIS. The critical point is that in a geometrical space of ducts, lobules, and acini, passive particle or cell distribution patterns may depend on the particles’ properties, including cell-cell cohesion and aggregate formation. In histopathology, this may appear to be an unconventional point of view. In other fields, potentially related phenomena are well-established. One possible example from geological sciences is the so-called granular segregation or hydraulic sorting. This process describes the differential transportation and deposition of riverbed sediments as a function of grain size, channel morphology, flow velocity, and many other factors (31,44–47).

The current interpretation of LCIS is “independent tumor initiation in multiple lobules at the same time,” based on tissue inspections from the 1940s (11). Still, recent studies indicate this concept is incorrect (8,20). Despite this discrepancy, it remains the prevailing theory due to the lack of alternative hypotheses. We propose an alternative “redistribution concept,” suggesting that loss of cell cohesion explains the distinct distribution of LCIS cells, supported by decades of evidence on cell adhesion loss in LCIS and invasive lobular carcinoma (starting with identifying CDH1/E-cadherin mutations) (4). Other, more directed cellular migration forces, such as chemotaxis to mammary lobules, could be a factor (48). We listed potential alternative candidate hypotheses in the Supplement (Supplemental Data Table 2) for an overview. Our modeling here only highlights that the redistribution hypothesis is a possible (and promising) candidate.

Formation of cohesive (or “sticky”) cell aggregates is typical for invasive breast carcinoma of no special type (NST-type) (49,50). These sticky cell aggregates render flow cytometry analyses of breast cancer cells more troublesome than in other tumor entities. Given that DCIS is a precursor of breast cancer (NST-type), it is reasonable to assume that DCIS cells form sticky aggregates, too. Despite the name “in situ,” the typical DCIS does not grow flat within the original ductal epithelium but within the ducts’ lumina. Presumably, early DCIS cells shed off from the epithelium and form cohesive aggregates within the open lumina of ducts. In this context, many morphologically different types of DCIS exist (51). The most common type of DCIS grows in solid cell aggregates or cohesive cell ribbons within the ducts’ lumina. Some rarer, particular kinds of DCIS do not grow as solid aggregates—the HMTR models LCIS with single beads and DCIS with bead aggregates, which is a simplistic approach that may still correctly reflect key molecular differences (non-cohesion versus cohesion).

We hope that our work stimulates new, much-needed research on LCIS. Studying LCIS is hampered by the lack of multiscale models (16,17). Computational and mathematical modeling can address the redistribution theory. The modeling presented here is a first step in this direction. It illustrates the concept but does not aim to provide an experimental or multi-scale theoretical environment that matches all physiologic conditions. It also does not address the potential impact of assortment in duct or lobules, as we assume a form of a well-mixed system. Beyond this assumption, future agent-based modeling will elucidate how well-mixedness impacts the modeling decisions. In this context, computational modeling of CIS cells may help elucidate LCIS distribution patterns, especially compared to DCIS. Computational modeling has recently explained comedo-type necrosis in DCIS (52,53). Other fields, such as granular segregation, could inform the modeling of LCIS (53). Taken together, our simplified experimental and mathematical modeling imply a potential mechanistic link between the loss of cell-cell cohesion and the predominantly lobular localization of LCIS. Our research may inspire further research on LCIS development and distribution.

## 5. Data availability

The data and code of this project are publicly available through the repository https://github.com/paltrock/LCIS_HTMR.

## 6. Acknowledgments

The authors thank Dipl.-Ing. J. Viering from the R&D technical workshop of the MHH for support with technical drawings and advice for the construction of the HMTR. The authors also thank Henriette Christgen for her technical assistance and permission to use the living room desk for the HMTR. RC, AT, and RC acknowledge generous funding from the Max Planck Society.

## 8. Statement of author contributions

All authors contributed to the study’s conceptualization, methods, formal analyses, writing, and manuscript editing. MC and ME contributed experimental components. MC designed the HMTR and executed the experimental work. RC and PMA developed the mathematical model. PMA performed statistical model fitting and parameter inference. MC and PMA wrote the manuscript with input from all authors.

