## Supplementary Tables and Figures for "Deficient cell-cell cohesion is linked with lobular localization in simplified models of lobular carcinoma *in situ* (LCIS)"

2: Departamento de Física, Universidade Federal do Paraná, Curitiba, Brazil

3: Clinic for Prosthetic Dentistry and Biomedical Material Science, Hannover Medical School, Germany

4: Department of Theoretical Biology, Max Planck Institute for Evolutionary Biology, Plön, Germany

5: MC and PMA contributed equally

Correspondence to:
Prof. Dr. med. Matthias Christgen, PhD
Institute of Pathology, Hannover Medical School
Carl-Neuberg-Str. 1, 30625 Hannover, Germany


Dr. rer. nat. Philipp Altrock
Department of Theoretical Biology, Max Planck Institute for Evolutionary Biology
August-Thienemann-Str. 2, 24306 Plön, Germany


### Supplemental Figure 1: constructional drawing of the HTMR

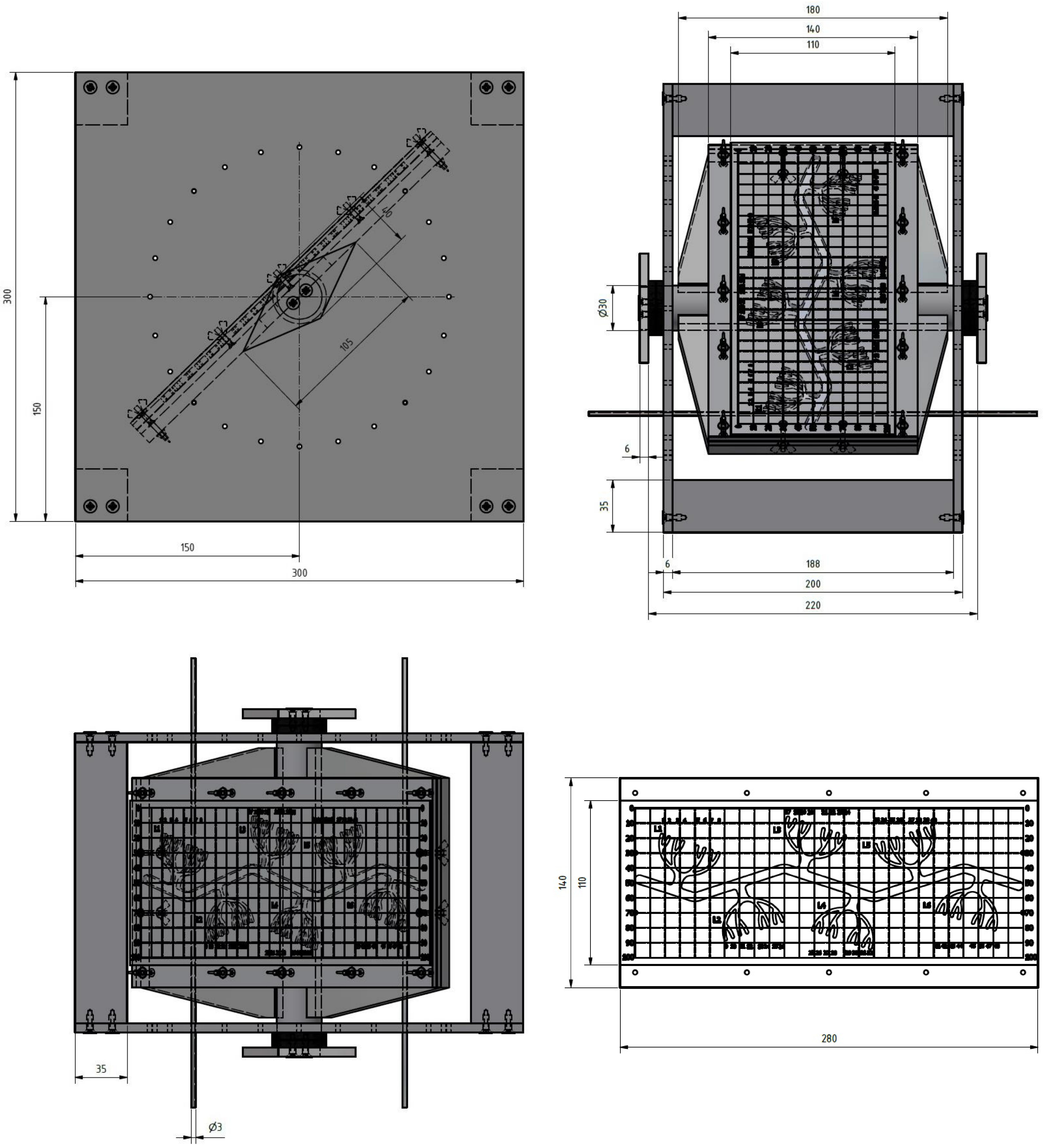


**Supplementary Figure 1.** HMTR, constructional drawing.

### Supplemental Figure 2: experimental procedure

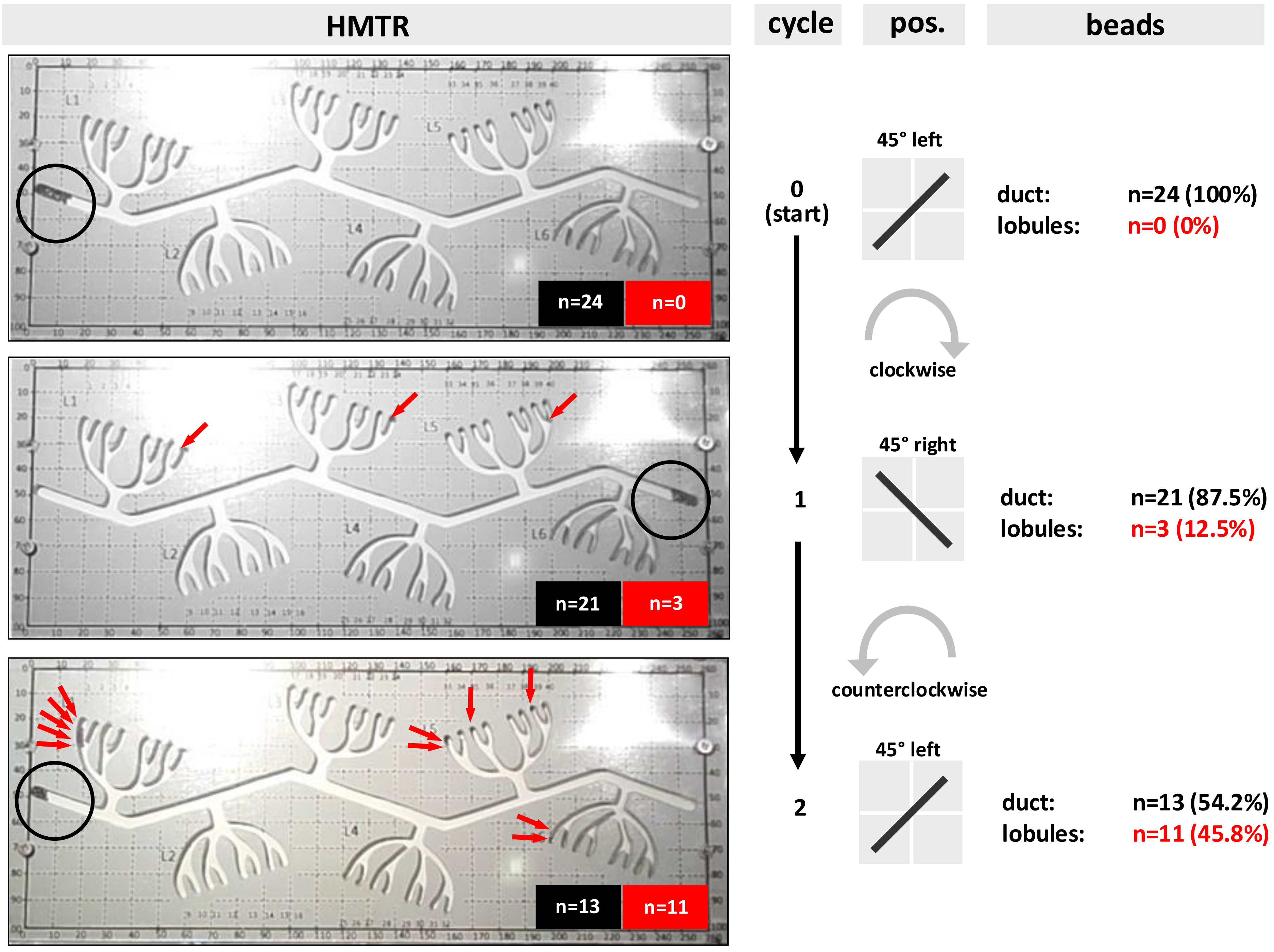


**Supplementary Figure 2.** Operation of the HMTR. From top to bottom, three panels illustrate two bead redistribution cycles. Top panel: Start position. Beads (n=24 single beads) were loaded in the left end of the duct. The rotary table rests on the left striker pin (45 degree to the left). Middle panel: Redistribution cycle 1. The rotary table was gently turned clockwise (45 degree to the right side). N=21 single beads moved to the right end of the duct, but n=3 beads (12.5%) entered lobules 1, 3 and 5 (red arrows). Lower panel: Redistribution cycle 2. The rotary table was gently turned counterclockwise (45 degree to the left side). N=13 single beads moved to the left end of the duct, but n=11 beads (45.8%) entered lobules 1, 5, and 6 (red arrows).

### Supplemental Table 1: Morphometric measurements in original histologic sections

Morphometric measurements

| | morphometric measurements | | | scaled 80:1 | | HM replica | |
|---|---|---|---|---|---|---|---|
| | n | median | range | median | range | actual sizes | actual/80 |
| terminal duct, diameter | 21 | 90 µm | 38 - 300 µm | 7.2 mm | 3.0 - 24.0 mm | 6.0 mm | 75 µm |
| acinus, outer diameter | 15 | 38 µm | 30 - 60 µm | 3.0 mm | 2.4 - 4.8 mm | - | - |
| acinus, inner diameter | 15 | 20 µm | 12 - 45 µm | 1.6 mm | 1.0 - 3.6 mm | 2.0 - 2.5 mm | 25 - 31 µm |
| lobule, diameter | 24 | 750 µm | 260 - 1400 µm | 60.0 mm | 20.8 - 112.0 mm | 40.0 mm | 500 µm |
| acini, number per lobule | 24 | n=31 | 10 - 98 | n=31 | 10 - 98 | n=8 | n=8 |

**Supplemental Table 2**

Overview of concepts and hypotheses for spatial LCIS development

| Hypothesis | Source | Key assumptions | Evidence for | | Evidence against | | Potential experimental validation strategies | Additional notes |
|---|---|---|---|---|---|---|---|---|
| | | | Authors | LOE | Authors | LOE | | |
| Initiation concept | Foote and Stewart, 1941 | LCIS resides in lobules because LCIS arises from lobuli. Multifocality of LCIS is viewed as independent tumor initiation in multiple lobules at the same time. | 1.) Foote and Stewart 1941, and traditional histopathology: Interpretation of histological sections. | LOE 5 (expert opinion) | 1. Wellings et al. 1976.: Interpretation of histological sections. | LOE 5 (expert opinion) | None. | Experimental validation is hampered by the lack o appropriate models. The biological differences of mammary lobules in mice and humans preclude experiments in laboratory animals (see introducti section of the manuscript). |
| | | | | | 2. Sakr et al. 2016 and Lee et al. 2019: Multifocal LCIS lesions harbor single unique somatic CDH1 mutations. | LOE 4 (cases series) | | |
| Redistribution concept | This work | Tumor initiation occurs anywhere in the epithelium. At the initiation site, LCIS cells shed off from the transformed epithelium. Subsequently, LCIS cells scatter through mammary ducts and colonize various lobules. | 1.) Christgen et al. 2024.: Interpretation of histological sections and the simplified HMTR model used for illustration. | LOE 5 (expert opinion and "first principle" research) | None. | n.a. | 1. Intra-ductal injection of LCIS cells transduced with appropriate reporter genes, followed by harvesting of the breast tissue for LCIS mapping years after the injection.<br>2. Computational modeling considering mammary gland anatomy, cell growth, cell death, cell adhesion, cell-cell interaction, tissue flexibility, intra-glandular fluid pressure *et cetera*.<br>3. Organoid models of realistic structures (Caruso et al. 2022 frontiers in Physiology 13:826107), yet to be combined with appropriate cancer cells. | 1. This theoretical validation strategy would be ur in humans. In laboratory animals, this theoretical approach would be hampered by the limited life s mice and the different biology of the mammary gl (non-permanent lobules).<br>2. This theoretical validation strategy is hamperec fact that the physical micro-environmental condit inside the mammary gland are not known.<br>3. The experimental implementation might be fea impacted by the ability to combine organoid deve and carcinogenesis. |
| Embryonic field cancerization concept | ELBCC members 2022; congress discussion | A heterozygous somatic CDH1 founder-mutation occurs in the mammary epithelium very early in life (e.g. during embryonic duct development). A mosaic of epithelial cells harboring the heterozygous CDH1 founder-mutation spreads over the mammary gland and causes LCIS later in life. | ELBCC 2022; congress discussion | below LOE 5 (expert opinion) | To our knowledge, there are no reported cases of LCIS and/or ILC in children. This may argue against this concept. | n.a. | Complete FFPE-embedding of mastectomy specimens from LCIS patients, followed by laser capture microdissection and NGS sequencing (CDH1) of the normal epithelium between LCIS foci. | High costs. Complete embedding of whole mastec specimens with 3D reconstruction of the branchir mammary system is extremely difficult and has sc been achieved only for few individual specimens ( Othake et al. 2001 Cancer 91:2263). Contaminatic by pagetoid LCIS cells is difficult to exclude. |
| Chemotaxis concept | This work. | LCIS cells grow in mammary lobules, because a chemotactic factor drives their motility towards lobules. | Still undetermined | below LOE 5 (expert opinion) | To our knowledge, no pro-lobular chemotactic factor is known, so far. | | Ex-vivo cultivation of human mammary lobules followed by Boyden chamber experiments for chemotaxis using LCIS cells or ILC cell lines. | Such an experiment would be complicated by the difficulties associated with the cultivation of hum: mammary epithelial cells in cell culture. Also, LCIS cannot be cultivated, while established ILC cell lin biased towards highly aggressive ILC variants hark additional TP53 mutations (Sflomos et al. 2021, C: 15:3299). |

**Abbreviations:**

ELBCC, European Lobular Breast Cancer Consortium; FFPE, formalin-fixed paraffin-embedded; ILC, invasive lobular carcinoma; LCIS, lobular carcinoma in situ; LOE, Oxford level of evidence (https://www.cebm.ox.ac.uk/resources/levels-of-evidence/ocebm-levels-of-evidence); NGS, next generation sequencing;
